# Similar neural pathways link psychological stress and brain health in health and multiple sclerosis

**DOI:** 10.1101/2022.12.19.521098

**Authors:** Marc-Andre Schulz, Stefan Hetzer, Fabian Eitel, Susanna Asseyer, Lil Meyer-Arndt, Tanja Schmitz-Hübsch, Judith Bellmann-Strobl, James H. Cole, Stefan M. Gold, Friedemann Paul, Kerstin Ritter, Martin Weygandt

**Affiliations:** Charité – Universitätsmedizin Berlin (corporate member of Freie Universität Berlin, Humboldt-Universität zu Berlin, and Berlin Institute of Health), Department of Psychiatry and Psychotherapy, Berlin, Germany; Bernstein Center for Computational Neuroscience, Berlin, Germany; Charité – Universitätsmedizin Berlin, corporate member of Freie Universität Berlin, Humboldt-Universität zu Berlin, and Berlin Institute of Health, Berlin Center for Advanced Neuroimaging, 10117 Berlin, Germany; Charité – Universitätsmedizin Berlin, corporate member of Freie Universität Berlin, Humboldt-Universität zu Berlin, and Berlin Institute of Health, NeuroCure Clinical Research Center, 10117 Berlin, Germany; Charité – Universitätsmedizin Berlin, corporate member of Freie Universität Berlin, Humboldt-Universität zu Berlin, and Berlin Institute of Health, Department of Neurology, 10117 Berlin, Germany; Charité – Universitätsmedizin Berlin, corporate member of Freie Universität Berlin, Humboldt-Universität zu Berlin, and Berlin Institute of Health, Regenerative Immunology and Aging, BIH Center for Regenerative Therapies, 13353 Berlin, Germany; Max Delbrück Center for Molecular Medicine and Charité – Universitätsmedizin Berlin, corporate member of Freie Universität Berlin, Humboldt-Universität zu Berlin, and Berlin Institute of Health, Experimental and Clinical Research Center, 13125 Berlin, Germany; Centre for Medical Image Computing, Department of Computer Science, University College London, London, UK; Dementia Research Centre, Institute of Neurology, University College London, London, UK; Institute of Neuroimmunology and Multiple Sclerosis (INIMS), Center for Molecular Neurobiology Hamburg, University Medical Center Hamburg-Eppendorf, 20251 Hamburg, Germany; Charité – Universitätsmedizin Berlin, corporate member of Freie Universität Berlin, Humboldt-Universität zu Berlin, and Berlin Institute of Health, Department of Psychiatry and Psychotherapy, Campus Benjamin Franklin, 12203 Berlin, Germany; Charité – Universitätsmedizin Berlin, corporate member of Freie Universität Berlin, Humboldt-Universität zu Berlin, and Berlin Institute of Health, Department of Psychosomatic Medicine, 10117 Berlin, Germany

**Keywords:** Multiple sclerosis, brain-age, psychological stress, functional connectivity, machine learning, convolutional neural networks.

## Abstract

Clinical and neuroscientific studies suggest a link between psychological stress and reduced brain health - in healthy humans and patients with neurological disorders. However, it is unclear which neural pathways mediate between stress and brain health and whether these pathways are similar in health and disease. Here, we applied an Arterial-Spin-Labeling MRI stress task in 42 healthy persons and 56 with multiple sclerosis. We tested whether brain-predicted age differences (“brain-PAD”), a highly sensitive structural brain health biomarker derived from machine learning, mirror functional connectivity between stress-responsive regions. We found that regional neural stress responsivity did not differ between groups. Although elevated brain-PAD indicated worse brain health in patients, anterior insula-occipital functional connectivity correlated with brain-PAD in both groups. Grey matter variations contributed similarly to brain-PAD in both groups. These findings suggest a generic connection between stress and brain health whose impact is amplified in multiple sclerosis by disease-specific vulnerability factors.

## Introduction

Converging evidence suggests that psychological stress can impair brain health – both under healthy conditions and in disease. For example, the application of stress-reducing meditation techniques is linked to better brain health in healthy humans (Luders et al., 2016). Work on multiple sclerosis (MS) showed that brain lesion activity in persons with MS (PwMS) increases during wartime (Yamout et al., 2010) and that therapeutic stress reduction can lower the occurrence of novel lesions in PwMS (Burns et al., 2014; Mohr et al., 2012). A large clinical cohort study showed that US veterans diagnosed with Post-Traumatic Stress Disorder (PTSD) have nearly double the risk of developing dementia over a seven-year period (Yaffe et al., 2010) and a meta-analysis that PTSD patients have reduced brain volume (Bromis et al., 2018). Consistently, stress – brain health associations were also found in invasive animal work. In healthy rodents, immobilization stress reduces hippocampal neurogenesis (Chetty et al., 2014), induces irreversible dendrite loss (Radley et al., 2006), and increases Amyloid-β peptides (Ray et al., 2011). The experience of early life traumata induces resistance to interferon-β and neurodegeneration mediated by lymphotoxin and chemokine receptors in an animal model of MS (Khaw et a., 2021).

Given these findings, one may ask whether a generic neurobiological mechanism exists that links psychological stress to reduced brain health independent of the presence of disease – a fact that could advocate the application of stress reduction for promoting brain health in health and disease alike. However, research that explicitly compares stress-brain health pathways, and that could potentially answer this question, is scarce. An explanation for this scarcity might be the presumption that non-invasive methods applicable in humans provide insufficient sensitivity – to measure neural stress processing, to identify meaningful brain health variations in healthy persons, or to separate disease-from age-related brain tissue variations. Neuroimaging, however, has shown that these conjectures are incorrect. Regarding the ability to measure neural stress processing, functional MRI (fMRI) has shown that psychological stress reliably correlates with specific patterns of regional brain activity (e.g., in prefrontal, limbic, insular, and occipital areas; Weygandt et al., 2016; Dedovic et al., 2009; Wang et al., 2005) and their functional connectivity (FC; e.g., Maron-Katz et al., 2016). Notably, FC might be of particular importance for studying stress processing in clinical contexts, as the impact of stress can be modulated by coping strategies (Folkman & Lazarus, 1988) and the neurobiological mechanism of coping (Ochsner et al., 2004) as well as the preference for specific coping strategies (i.e., benecifial or detrimental for well-being) depend on FC (Santarnecchi et al., 2018). Keeping this in mind, the specific suitability of FC for studying neurobiological stress-brain health pathways is underscored by the findings of Burns et al. (2014), who showed that the impact of stressful life events on brain lesion activity was modulated by the MS patients’ subjective evaluation of (i.e., coping with) life events as either positive or negative. With respect to the ability of measuring brain health in a non-invasive fashion, the “brain-age” paradigm which assesses brain health via “brain-predicted age differences” (“brain-PAD”; Cole et al., 2020) has proven to fulfill the rigorous sensitivity requirements for detecting not only disease-related tissue alterations but also subclinical variations in healthy persons (HPs). Specifically, brain-PAD measures deviations from normal brain aging in terms of a formula that could be expressed as “a person’s age predicted by a machine learning algorithm (i.e., their “brain-age”) through a comparison of their anatomical MRI to those of a healthy reference cohort covering varying ages minus their chronological age”. Algorithms successfully trained to predict the chronological age of HPs overestimate it (i.e., compute a positive brain-PAD) for persons with MS, dementia, mild-cognitive impairment, and schizophrenia (Kaufmann et al., 2019). In MS, this has been underlined by a MAGNIMS consortium study (Cole et al., 2020), which additionally shows that brain-PAD is also a biomarker for disease course prediction as higher brain-PAD goes along with shorter time-to-disability progression. Moreover, brain-PAD has sufcicient sensitivity to detect meaningful brain health variations in HPs as brain-PAD links positively to mortality risk in healthy elderly (Cole et al., 2018) and negatively to meditation practice in HPs (Luders et al., 2016). Finally, with respect to age-related brain tissue variations, the method intrinsically controls for the impact of age on brain health given the abovementioned subtraction approach.

Despite the high individual sensitivity of FC and brain-PAD, however, these techniques have not yet been combined to identify and compare poorly understood stress-brain health pathways in HPs and neurological patients. Therefore, we employed a well-established Arterial-Spin-Labeling (ASL) fMRI stress paradigm comprising a rest and a socially evaluated mental arithmetic stress condition (e.g., Brasanac et al., 2022; Meyer-Arndt et al., 2021; 2020; Weygandt et al., 2016; Wang et al., 2007; 2005) to compare neurobiological pathways linking stress and brain health in 42 HPs and 56 PwMS. PwMS represent a group of patients well able of dealing with experimental tasks given their comparably moderate average disability and age (Manouchehrinia et al., 2017). ASL is a quantitative MRI method reflecting functional processes via differences in the regional cerebral blood flow (rCBF; ml/100g/min) between experimental conditions that is increasingly used for FC characterization. It is better suited for stress measurement than the alternative technique blood-oxygenation-level-dependent (BOLD) fMRI as rCBF is more robust towards temporally slow (Wang et al., 2005; 2003; Aguirre et al., 2002) and thus stress-like (Kirschbaum et al., 1993) signal artifacts. The review by Chen et al. (2015) and the Discussion provide further details on ASL-based FC.

We conducted four main analyses. The first three were cross-sectional and based on one set of participants, the fourth was longitudinal and based on a subset. In the first main analysis, we evaluated *activity* differences of regions defined in a brain atlas as indicators of regional stress responsivity. Specifically, we computed the regions’ spatially averaged voxel rCBF timeseries for the resting and the stress condition in each HP and each PwMS and contrasted the conditions’ timeseries within and between groups. We expected that regional stress responsivity would not differ between groups. In the second main analysis, we computed the *functional connectivity* for stress-responsive regions and tested associations with brain-PAD. In particular, we computed the correlation between spatially averaged rCBF timeseries for each pair of stress-responsive regions separately for the stress and rest condition in each participant and related these parameters to brain-PAD across participants separately for each condition and group. We expected that stress-stage FC of a similar pair of regions would be associated with brain-PAD in both groups. In the third analysis, we evaluated associations between regional brain aging and whole-brain grey matter (GM) fraction in PwMS and HPs to test our assumption that brain-age, the endpoint of the presumed stress-brain health pathway, has a comparable neurobiological substrate in both groups. Finally, in the fourth main analysis, we tested associations between resting and stress-stage FC assessed at the fMRI visit of our study and future brain-PAD variations in PwMS occurring in a follow-up period and expected that stress-stage FC would contain prognostic information for future brain health.

## Materials and Methods

### Participants

The study integrates data from two study projects conducted by the NeuroCure and the Experimental and Clinical Research Center at Charité – Universitätsmedizin Berlin, Germany. Stress fMRI data acquired in the first project were previously presented in Weygandt et al. (2016) and Meyer-Arndt et al. (2021; 2020), those in the second in Brasanac et al. (2022). Patient inclusion criteria were comparable across projects: Meeting the 2010 McDonald Criteria (Polman et al., 2011) for relapsing-remitting or secondary-progressive MS (first project) or those for relapsing-remitting MS (second project). Patients had to receive a stable disease-modifying treatment for at least six (first project) or three (second project) months or no disease-modifying treatment in these periods. Finally, they had to be ≥ 18 years of age and the capabilities to use the experimental devices without restrictions. In both projects, patients were excluded if they had an additional neurologic disorder, MRI contraindications including pregnancy, an acute relapse, received steroid treatment in the last four weeks, or in case of insufficient anatomical image quality (visually assessed by M.W.). Moreover, patients were excluded in case of a known psychiatric diagnosis (first project) or when a psychiatric disorder other than a depressive or anxiety disorder was diagnosed by an experienced psychotherapist (second project). If applicable, these inclusion and exclusion criteria were identical for HPs in both projects. To determine the participants of main analyses 1 - 3, we then proceeded by excluding three participants from the 57 participants in Weygandt et al. (2016) and seven of 66 participants in Brasanac et al. (2022) who passed the abovementioned criteria but showed pronounced fMRI head motion as indicated by the Framewise Displacement quality assurance metric (Power et al., 2014; see Supplement for details). Prior to data pooling, we finally excluded the data acquired in the first project (the project with slightly lower anatomical MRI resolution) of those 15 persons who participated in both projects to avoid mixing up within- and between-subject variation. Consequently, data of 39 / 59 participants were entered in the three main analyses from the first / second project to a total of 98 participants (56 PwMS, 42 HPs). To determine the participants of main analysis 4 exclusively taken from the first study project, we excluded a single participant from the 25 in Meyer-Arndt et al. (2020) who passed the abovementioned criteria, but showed pronounced fMRI head motion. For the 24 remaining PwMS, the median delay between the first (acquisition of fMRI data and first anatomical MRI session) and the follow-up visit (second anatomical MRI session) was 1017 (range 717 – 1439) days. Both projects were conducted in accordance the Helsinki Declaration of 1975 and approved by the ethics committee of Charité – Universitätsmedizin Berlin (first project: EA1/182/10, amendment V; second: EA1/208/16). Written informed consent was obtained from all participants at all time points.

### Data Availability

Anatomical MRI images will not be made available due to privacy issues of clinical data. Access to data other data will be granted by the corresponding author on request depending on the approval by the ethics committee and under a formal Data Sharing Agreement.

### Clinical assessments

The EDSS (Kurtzke, 1983) was used to evaluate clinical disability. In the first project, BDI-II (Hautzinger et al., 2009) was used to measure the severity of depressive symptoms, BDI-I (Beck et al., 1961) in the second. We used the algorithm described by Wahl et al. (2014) to convert the scores obtained with both measures into one common metric. For other measures assessed, see Weygandt et al. (2018).

### fMRI stress paradigm

In project one, the paradigm comprised seven stages (1. Rating I, 2. Rest, 3. Rating II, 4. Stress, 5. Rating III, 6. Rest II, 7. Rating IV), in project two the first five thereof were assessed. Stress stage 4, during which participants had to conduct a series of mental arithmetic tasks, was divided in “Evaluation” (4a) and “Feedback” (4b) substages. In the present study, we evaluated the data from the first five stages consistently acquired in both projects. Perceived psychological stress was measured in the 1^st^, 3^rd^, and 5^th^ stage (2 min duration each) via a nine-point Likert-scale ranging from “not at all” to “strongly”. ASL scans and heart rate (Supplement) were acquired in the 2^nd^ (8 min) and 4th (12 min) stage. Fig. 1 gives details.

**Figure 1.**
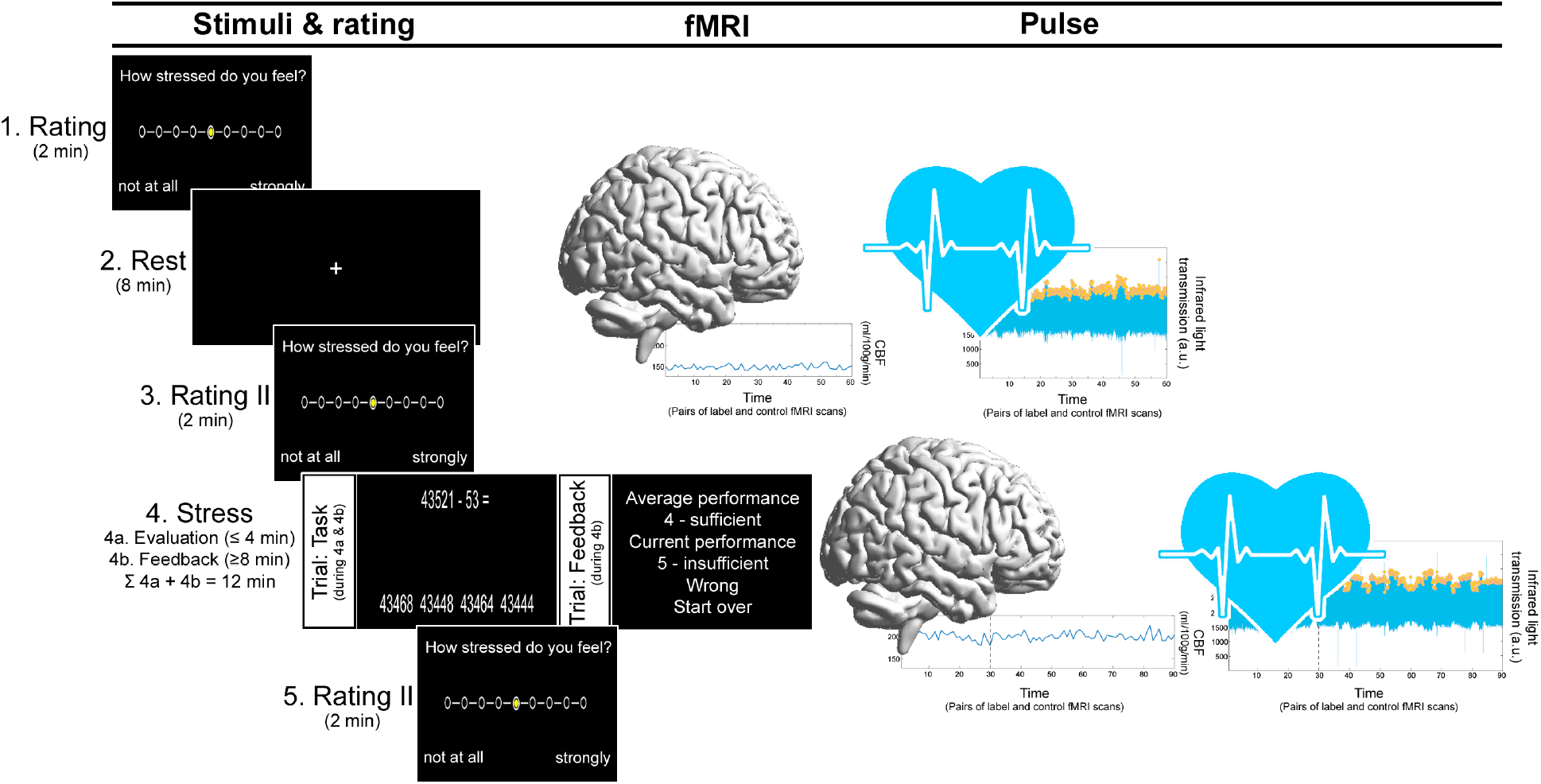
fMRI Stress paradigm. The total paradigm comprised five stages. The stress stage was divided in 4a (“Evaluation”) and 4b (“Feedback”). In 4a, participants performed subtraction tasks across a series of trials. In each, they had to subtract operand Y from X as fast as possible by selecting the correct answer among a set of four options depicted on a screen with an MRI-compatible response box. X started with 43521. In all trials in 4a (and 4b), operand Y was randomly determined (range: 1 – 99). In the first project, 4a either lasted until a participant provided ten correct answers or 4 minutes if they failed to provide ten correct answers. In the second, 4a had a fixed duration of 4 minutes. To compensate, we only evaluated rCBF, heart rate, and mental arithmetic performance data acquired in the final 8 minutes of 4b and modelled the duration of 4a as CNI “time-to-feedback” in respective regression analyses in this study. In 4a, the time provided to solve a task was 8 seconds. When a trial was correctly solved, X equaled the difference X minus Y from the preceding trial. X remained unchanged in case of a false or too slow answer. After 4a, 4b started immediately. 4b differed in three points. First, feedback (i.e., German school grades ranging from “1 – very good” to “5 – insufficient”) was provided in each trial. The grade was determined by the difference between the participant’s fastest correct response in 4a and the response time of a given trial in 4b. For false or too slow answers, the feedback was always “5 – insufficient”. In this case, second, X was reset to 43521. Finally, third, the time for response selection was adapted to the participant’s performance. Starting with 8 seconds at the beginning of 4b, this time was reduced by 10% when a correct answer was provided and increased by 10% in case of false/too slow answers. This adaptive mechanism was implemented to ensure that each participant operated at the individual performance maximum across the paradigm and was thus comparable across participants from the perspective of a subjective performance norm.

### MRI acquisition

All MR images were acquired with the same 3 Tesla whole-body tomograph (Magnetom Trio, Siemens, Erlangen, Germany) and standard 12-channel head coil. Two anatomical MR sequences were acquired in both projects. In the first, a T1-weighted sagittal 3D MP-RAGE sequence (176 slices; slice thickness 1.3 mm; in-plane voxel resolution 1.5 · 1.5 mm^2^; TR = 1720ms; TE = 2.34ms; FA = 9°; FOV = 192 · 192 mm^2^; matrix size = 128 × 128; duration 1 min and 43 sec) and a sagittal T2-weighted sequence were acquired (176 slices; 1 mm isotropic voxels; TR = 5000 ms; TE = 502 ms; FA = 120°; FOV = 256 · 256 mm^2^; matrix size = 256 · 256; 5 min and 52 sec duration). In the second, a sagittal 3D MP-RAGE sequence (176 slices; 1 mm isotropic voxels; TR = 1900 ms; TE = 3.03 ms; FA = 9°; FOV = 256 · 256 mm^2^; matrix size = 256 × 256; 4 min 26 sec duration) and a sagittal T2-weighted FLAIR sequence (176 slices; 1 mm isotropic voxels; TR = 6000 ms; TE = 388 ms; TI = 2100 ms; FA = 120°; FOV = 256 · 256 mm^2^; matrix size = 256 · 256; 7 min 44 sec duration). In both projects, perfusion brain images were acquired with the same pseudo-continuous ASL (pCASL) echo-planar imaging (EPI) sequence (Wang et al., 2005) comprising 22 ascending transversal slices covering the whole brain (slice thickness 5.75 mm; 15% inter-slice gap; in-plane voxel resolution 3 · 3 mm^2^; TR = 4000 ms; TE = 19ms; FA = 90°; FOV = 192 · 192 mm^2^; matrix size = 64 · 64; label duration 1.5 sec, post-label delay 1.2 s; phase-encoding direction anterior to posterior). Two spin-echo EPI reference volumes with opposing phase encoding directions (anterior to posterior, posterior to anterior) were acquired prior to the ASL measurements with matching readout and geometry for distortion correction.

### MRI preprocessing

#### Anatomical scans

First, a manual lesion mapping using T2-weighted scans was conducted. Standard preprocessing performed with SPM12 (Wellcome Trust Centre for Neuroimaging, Institute of Neurology, UCL, London UK, http://www.cil.ion.ucl.ac.uk/spm) comprised a spatial mapping of participants’ T2-weighted to T1-weighted scans, a spatial normalization of T1-weighted scans to the Montreal Neurological Institute (MNI; Tzourio-Mazoyer et al., 2002) standard space and a segmentation of T1-weighted images into GM, white matter (WM) and cerebro-spinal fluid. Segmented tissue voxel maps were used for computation of tissue specific group-masks and whole-brain GM fraction. See Supplement for details.

#### Functional scans

Preprocessing comprised head motion correction, B° distortion correction, coregistration to the MNI standard space (voxel resolution 3 · 3 · 3 mm^3^) utilizing the spatial normalization parameters computed for the anatomical images, and spatial smoothing. Subsequently, we extracted voxel-wise rCBF timeseries for all participants and rest and stress separately. Preprocessing and all subsequent steps utilized data from the full eight minutes of the rest and the final eight minutes of the stress condition. We focused on the final eight minutes of the stress condition as these were consistently included in the “Feedback” stress-substage in all participants. For details, see Supplement.

### Brain-age, brain-PAD, and voxel maps of regional brain-age

To determine the participants’ brain-age, their T1-weighted scans normalized to MNI space were skull stripped to prepare these images for brain-age computation first. In the next step, we used a convolutional neural network (CNN) provided by Bashyam et al. (2020) for brain-age prediction which was pre-trained for this purpose based on the anatomical MRI scans of 11,729 healthy persons. The CNN analyzes the individual MRI slices of a participant’s brain scan and predicts the participant’s age from each slice. As in Bashyam et al. (2020), the median age predicted across all slices was used as a measure of brain age. The difference “predicted brain-age minus chronological age” served as brain-PAD marker in the cross-sectional analyses. The difference in brain-PAD computed at follow-up minus at baseline (Δbrain-PAD) served as marker of future brain-PAD variations in the longitudinal analysis. To determine one map per participant reflecting the participants’ regional brain aging for main analysis three, we entered the normalized T1-weighted scans into the Shapley Additive exPlanations toolbox (Lundberg & Lee, 2017) and computed voxel-wise brain-age scores. In these voxel maps, positive scores bias a person’s the brain-age towards an older age, negative towards a younger one. The resulting maps were smoothed with a 3D gaussian kernel (eight-millimeter full width at half maximum).

### Within-participant regional stress responsivity

To determine the regional neural stress responsivity, we first computed an rCBF timeseries for each participant, condition, and each of the GM regions in the Neuromorphometrics brain atlas (http://Neuromorphometrics.com). Timeseries were averaged across activity of all voxels in a given atlas region that were covered by a group-specific GM mask and contained non-zero rCBF. If a region did not include a single voxel fulfilling these criteria in one or more participants of a group, this region was excluded from the group’s analysis. 121 of 122 atlas regions were included for HPs, 119 for PwMS. An explanation for this coverage pattern is provided in the Supplement. Regional within-participant stress-rest activity differences were computed as stress responsivity indicators by contrasting the regions’ averaged timeseries of both conditions via linear regression. The resulting regression coefficients were entered into main analysis 1. See Supplement for details.

### Functional connectivity modeling

In the next step, we computed the resting and stress-stage FC for each participant and all pairs of stress-responsive regions (indicated by main analysis 1). Specifically, the (Fisher Z-transformed) correlation coefcicient for average rCBF timeseries of a given pair of regions was computed for the resting and the stress-stage as FC measures. These pair-of-region-wise markers were entered into main analysis 2.

### Statistical analyses

#### Main Analysis 1: Regional neural stress response activity

We tested regional stress-rest brain activity differences within and between groups with robust linear regression with the abovementioned regression coefficients mentioned in section “Within-participant regional stress responsivity” above as dependent variable. Age, sex, study project, and depressive symptom severity (collectively referred to as “standard nuisance factors” in the following), cognitive task load (i.e., the average duration of the inter-trial interval in the final eight minutes in the stress condition) and time-to-feedback (the duration of the “Evaluation” stress substage) were included as covariates of no interest (CNI) in both within-group analyses. In the patients’ analysis, EDSS, disease duration, and the presence of a progressive disease form (y/n) were included as additional CNI. Permutation testing (10,000 permutations) was used for inference. For the patients’ and between-groups analyses, a family-wise-error (FWE) corrected significance threshold for undirected tests of α_FWE_ = 0.05 was computed by dividing 0.05 (threshold for single test) by 119 (the number of regions consistently included in the patients’ and the HPs’ analysis) and thus equaled 4.2 ∙ 10^−4^ on a single test level. For HP, the threshold corresponded to 0.05 / 121 = 4.1 ∙ 10^−4^. Here and for all other regression-based analyses, we report effect size measures f^2^ (f^2^ ≥ 0.02 weak, f^2^ ≥ 0.15 medium, and f^2^ ≥ 0.25 strong effect; Cohen, 1988). Analyses testing psychological (self-report data) and physiological stress responses (heart rate) are described in Supplementary analysis 1.

#### Main Analysis 2: Functional connectivity and brain-PAD

We used robust linear models to regress participants’ brain-PAD on resting and stress-stage FC between pairs of stress-responsive regions independently for both groups and conditions. CNI were the same as in main analysis 1. Permutation testing (10,000 permutations) and an FWE-corrected significance threshold were used to evaluate significance in undirected pair-of-region-wise tests. We defined an α_FWE_ of 0.05, which corresponded to 0.05 / (17 · [17 – 1] / 2) = 3.68 · 10^−4^ on a single test level for HPs because 17 regions with significant stress-responses were found in HPs (see Results). Since 14 such regions were found in PwMS, the corresponding threshold for PwMS was 0.05 / (14 · [14 – 1] / 2) = 5.49 · 10^−4^. We conducted two related supplementary analyses. In the first (Supplementary analysis 2), we determined FC - grey matter fraction associations to compare them against FC – brain-PAD associations or to evaluate the relative suitability of brain-PAD for studying stress – brain health pathways respectively. In the second (Supplementary analysis 3), associations between regional neural stress response activity and brain-PAD were evaluated to test the relative suitability of FC.

#### Main Analysis 3: Group similarities and differences in regional brain aging

To clarify whether brain-age, the presumed endpoint of the investigated stress-brain health pathway, has comparable neurobiological substrates in both groups, we tested associations between regional brain-age and the whole-brain GM fraction in both groups separately. Additionally, we tested associations between regional brain aging and whole-brain T2-weighted lesion load in patients and differences in regional brain aging between groups to further evaluate potential pathobiological contributors to regional brain-age. For this purpose, the regional brain age maps were analyzed with SnPM13 (https://warwick.ac.uk/fac/sci/statistics/staff/academicresearch/nichols/software/snpm/), an SPM12 toolbox using permutation testing (10,000 permutations) and the maximum statistic FWE-correction procedure for inference. In the within-group analyses (GM-fraction in both groups, lesion load in patients), the full set of covariates comprised the standard nuisance factors, information processing capacity (i.e., the number of correctly solved arithmetic tasks in the final eight minutes in the stress condition, a parameter reciprocal to cognitive task load due to the performance-adaptive task pace), GM and WM fraction for both groups. T2-weighted lesions, EDSS, disease duration, and progressive MS were additionally modeled in PwMS. Based on the full covariate set, one analysis was conducted testing associations between brain-age voxel scores as dependent variable and GM fraction as covariate of interest in both groups separately. All other covariates served as CNI. The same was done for lesion load in patients. In the analysis of group differences, standard nuisance factors plus information processing capacity served as CNI. All analyses except for the GM fraction analyses were constrained to areas located in the intersection of the SPM12’s brain mask and that of Bashyam et al. (2020); the former was additionally constrained to coordinates in a given group’s GM mask. Because higher GM fraction indicates better brain health but higher brain-age and lesion load indicate worse brain health, we tested for negative associations between GM fraction and brain-age voxel scores and for positive associations regarding lesion load (α_FWE_ = 0.05).

#### Main Analysis 4: Functional connectivity and future brain-PAD in PwMS

We evaluated whether rest and stress-stage FC between the pairs of stress responsive regions determined for PwMS is associated with future variations in brain-PAD (i.e., Δbrain-PAD). The regression model was identical to that in the second main analysis except for the facts that (i) Δbrain-PAD served as dependent variable, and (ii) brain-PAD at baseline, and (iii) the follow-up duration were modelled as additional CNI. We applied the significance threshold of α_FWE_ of 0.05 also applied in main analysis 2 for PwMS which corresponded to 0.05 / (14 · [14 – 1] / 2) = 5.49 · 10^−4^ on a single test level.

## Results

### Clinical and demographic participant characteristics

Ten of 56 patients in the cross-sectional data set were treated with β-interferons, ten with dimethyl fumarate, seven with glatiramer acetate, five with teriflunomide, and nine with fingolimod. Among the 24 patients in the longitudinal data set, four were treated at baseline with β-interferons, five with dimethyl fumarate, four with glatiramer acetate, one with teriflunomide, and five with fingolimod. Tab. 1 gives details.

**Table 1.** Demographic, clinical and neuroradiographic participant characteristics. Abbreviations: #, number; corr., correct; DD, disease duration assessed from the onset of symptoms; fract., fraction; Inf. proc., information processing capacity assessed in terms of the number of correctly solved mental arithmetic trials in the final 8 minutes of the stress stage; MD, median; PROG, denotes the presence of a progressive form of the disease; pts., points; RG, range; RR, relapsing remitting; SP, secondary progressive; T2LL, T2-weighted lesion load.

| Cross-sectional data set |  |  |  |  |  | Longitudinal data set |  |
| --- | --- | --- | --- | --- | --- | --- | --- |
|  |  | Persons with MS | Healthy persons |  |  |  | Persons with MS |
| **Sex**<br>(f/m) | # | 37 / 19 | 28 / 14 | $\chi^2$<br>**p** | 0.00<br>0.951 | # | 13/11 |
| **Age**<br>(yrs) | MN<br>SD<br>RG | 46.4<br>10.7<br>22 - 66 | 44.7<br>15.5<br>21 - 75 | **t**<br>**p** | 0.63<br>0.530 | MN<br>SD<br>RG | 48.3<br>7.6<br>27 - 62 |
| **Inf. proc.**<br>(#corr. trials) | MN<br>SD | 28.1<br>10.9 | 34.1<br>12.0 | **t**<br>**p** | -2.60<br>0.011 | MN<br>SD | 30.7<br>11.7 |
| **BDI**<br>(pts.) | MN<br>SD | 8.3<br>7.4 | 2.6<br>3.0 | **t**<br>**p** | 4.69<br>$9.1 \cdot 10^{-6}$ | MN<br>SD | 7.1<br>5.5 |
| **GM**<br>(fract.) | MN<br>SD | 0.38<br>0.05 | 0.41<br>0.05 | **t**<br>**p** | -3.07<br>0.003 | MN<br>SD | 0.41<br>0.04 |
| **WM**<br>(fract.) | MN<br>SD | 0.23<br>0.02 | 0.24<br>0.02 | **t**<br>**p** | -2.08<br>0.040 | MN<br>SD | 0.25<br>0.02 |
| **T2LL**<br>(cm<sup>3</sup>) | MD<br>RG | 931.5<br>13.5 -<br>$1.3 \cdot 10^4$ | | | | MD<br>RG | 730.7<br>13.5 -<br>$9.1 \cdot 10^3$ |
| **EDSS**<br>(pts.) | MD<br>RG | 2.5<br>0 - 6 |  |  |  | MD<br>RG | 3.5<br>1 - 6 |
| **DD**<br>(yrs) | MN<br>SD | 12.7<br>8.8 |  |  |  | MN<br>SD | 11.1<br>7.7 |
| **PROG**<br>(RR/<br>SP) | # | 49/7 |  |  |  | # | 18/6 |

### Brain-age prediction accuracy and group differences in brain-PAD

The correlation between chronological and brain-age in HPs was r = 0.90 (r^2^ = 0.81, p = 5.4 · 10^−16^; mean absolute error = 6.18 [SD = 4.83] years). The average brain-PAD was 10.08 (SD = 8.99) years in patients and 3.95 (SD = 6.82) years in HPs and thus significantly higher in PwMS (t = 3.69, p = 3.7·10^−4^).

### Main Analysis 1: Regional neural stress response activity

A wide-spread set of stress-responsive regions spanning frontal, temporal, parietal, occipital, and cerebellar areas was identified in both groups (i.e., 17 regions in HPs, 14 in PwMS). No group differences were found (Fig. 2).

**Figure 2.**
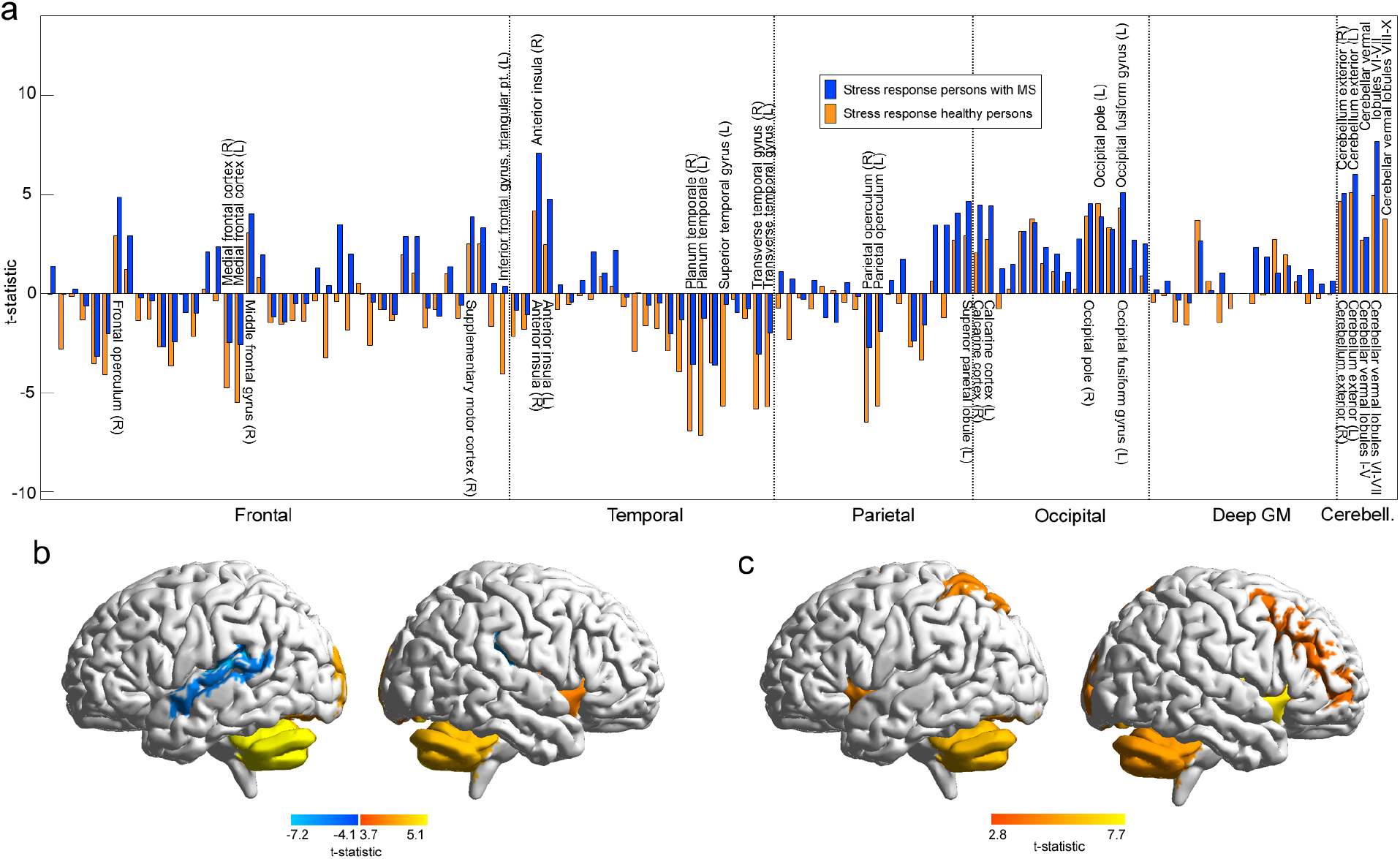
Regional neural stress response activity. 2a shows neural stress responses. The bar graph in 2a shows the t-statistics for the difference in rCBF for stress (final 8 minutes of stage 4b) minus rest (stage 2) separately for both groups. Labels depicted above the baseline of 2a highlight the 17 regions yielding significant rCBF differences in HPs, labels depicted below the 14 in PwMS. Please note, that the coarse clustering of regions in 2a into frontal, temporal, etc. was conducted to increase the comprehensibility of the graph and is not necessarily optimal in anatomical terms. The render brains in 2b (HPs) and 2c (PwMS) summarize regions with significant stress responses.

### Main Analysis 2: Functional connectivity and brain-PAD

We observed a significant positive association for stress-stage FC between right anterior insula and left occipital pole and brain-PAD in HPs (t = 4.22, p_uncorr._ = 4.0·10^−5^ = p_FWE_ = 0.005, f^2^ = 0.57). In PwMS, a significant positive association for FC during stress between left anterior insula and occipital fusiform gyrus during and brain-PAD was found (t = 3.86, p_uncorr._ = 1.3·10^−4^ = p_FWE_ = 0.012, f^2^ = 0.74). Resting-stage FC was neither related to brain-PAD in HPs, nor in PwMS (Fig. 3).

**Figure 3.**
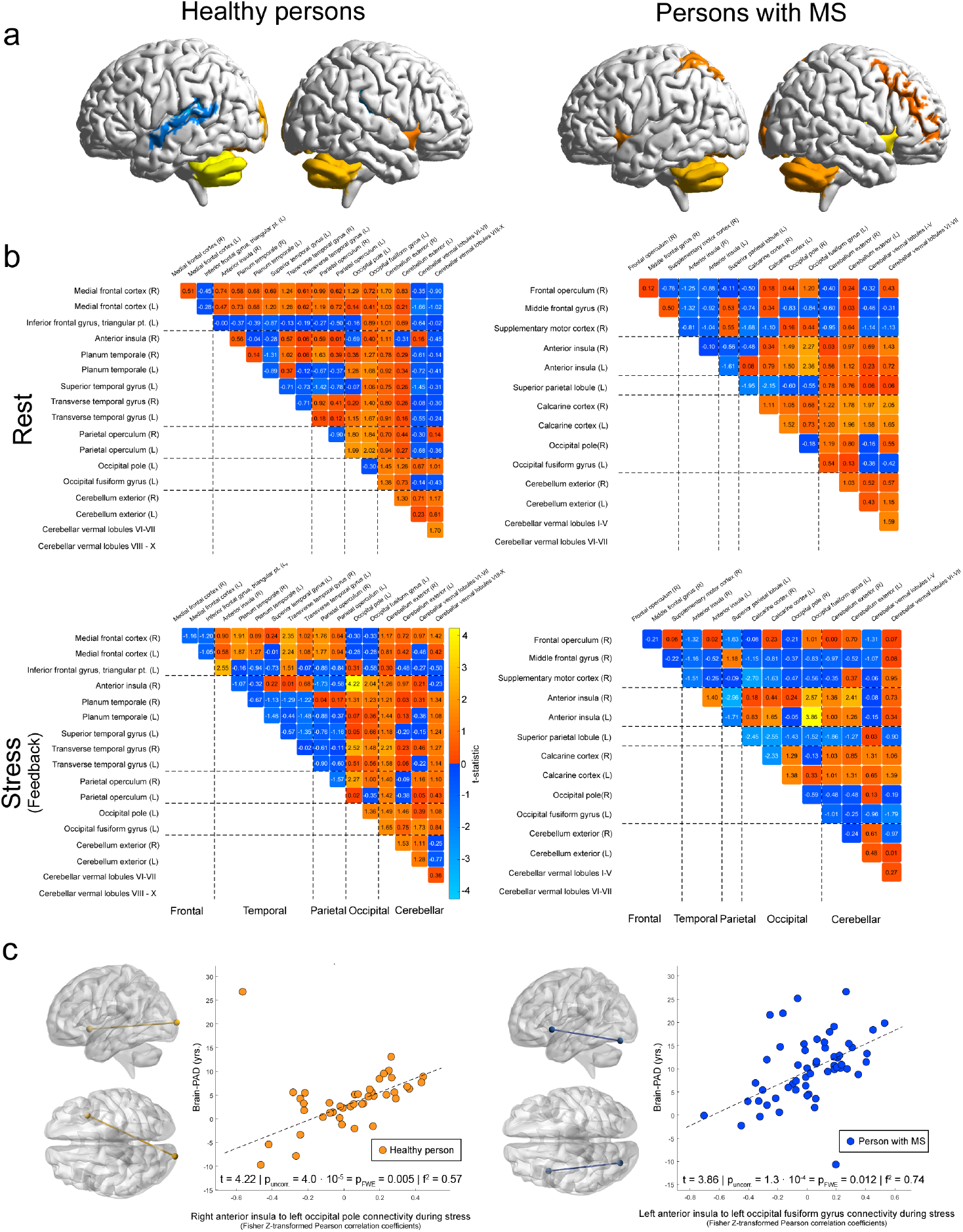
Functional connectivity and brain-PAD. 3a yet again depicts the stress-responsive regions (also shown in Fig. 2b, c) to ease readability of the figure. The heatmaps in 3b depict the t-statistics for the association between FC of a given pair of stress-responsive regions on one hand and brain-PAD on the other. Specifically, the left half of 3b shows these t-statistics for HPs, the right half for PwMS. The top row shows associations for rest, the bottom row for those for stress. In the left half of 3c, we illustrate (from left to right) the connection between (the center coordinates) of anterior insula and occipital regions predictive of brain-PAD and a scatter-diagram of the association between stress-related anterior insula – occipital FC and brain-PAD in HPs. In the right half of 3c, we depict the same for PwMS.

### Main Analysis 3: Group similarities and differences in regional brain aging

A strong negative association between brain-age voxel markers in parietal cortex and GM fraction was found in both groups. For HPs, the peak was located in left supramarginal gyrus at coordinate MNI: −54, −40, 38 (t = −6.25, p_FWE_ = 0.012, f^2^ = 1.15). In other words, the higher the regional brain-age of this coordinate in a HP (the higher this coordinate biased the CNN to predict an older brain-age), the lower the overall GM fraction for this HP. In PwMS, the peak coordinate was located directly adjacent in angular gyrus (MNI: −48, −57, 41; t = −8.85, p_FWE_ = 0.0002, f^2^ = 1.78). A significant negative association in angular gyrus regions that overlapped in both groups was found at MNI: −47, −51, 44 (HPs: t = −5.75, p_FWE_ = 0.041, f^2^ = 0.97; PwMS: t = −6.00, p_FWE_ = 0.008, f^2^ = 0.82). Moreover, a strong positive association of regional brain aging with whole-brain lesion volume in periventricular white matter (WM; MNI: 33, −49, 10; t = 8.45, p_FWE_ = 0.0002, f^2^ = 1.62) was found in PwMS. Finally, the analysis testing for group differences showed that PwMS were characterized by higher brain age of right thalamus proper than HPs (MNI: 18, −27, 11; t = 5.51, p_FWE_ = 0.0038, f^2^ = 0.33). For an illustration see Fig. 4, for a complete list of coordinates identified in regional brain aging analyses, see supplementary Tab. S1.

**Figure 4.**
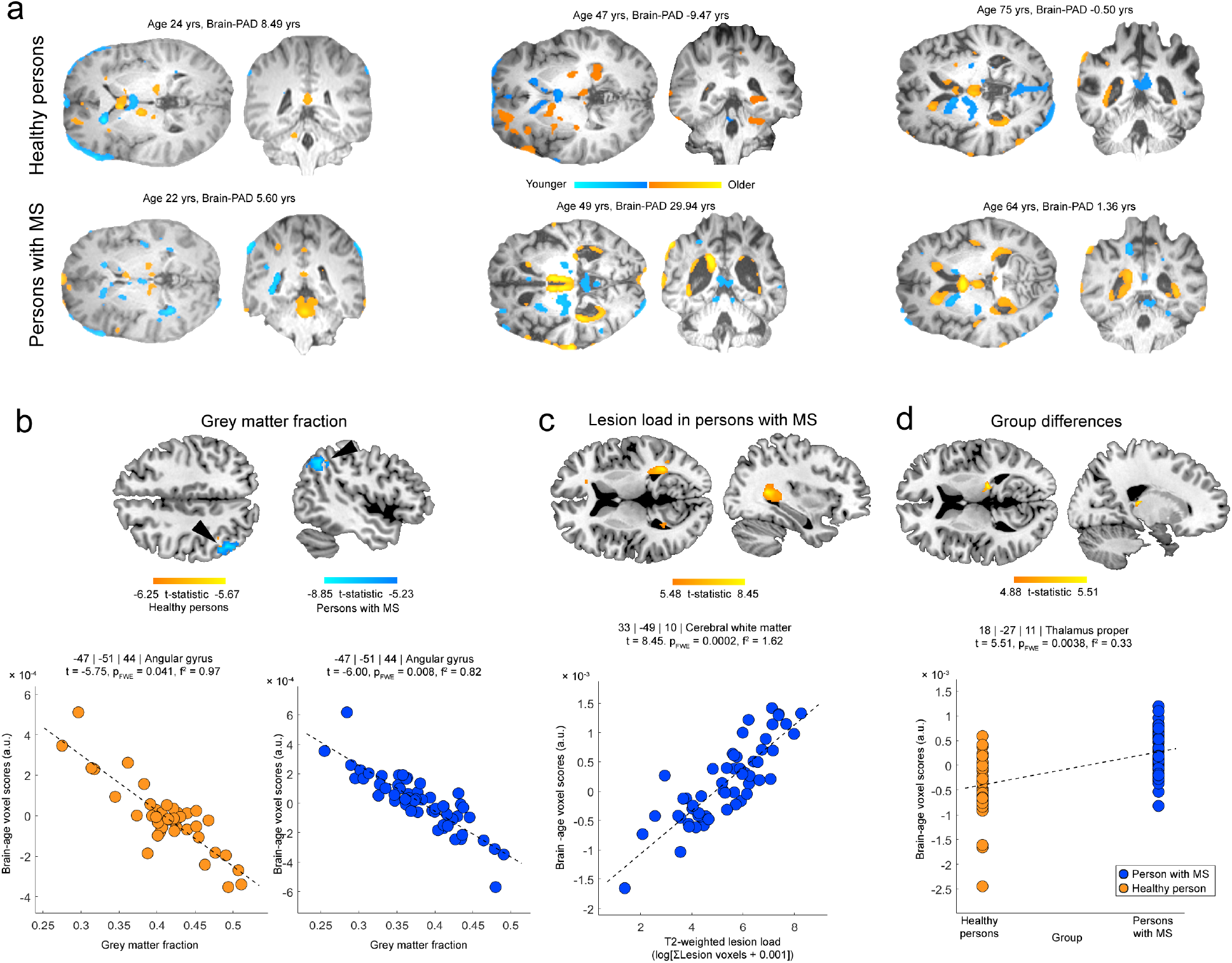
Group similarities and differences in regional brain aging. 4a first illustrates the brain age prediction approach or how the machine learning algorithm predicts the brain age for six selected participants respectively. Specifically, it determines a score for each voxel which individually biases the algorithm slightly to a younger or an older age and which can be understood as marker of regional brain aging. The (weighted) sum of voxel scores gives the brain age of a person. Voxels highlighted in blue correspond to the 5% of all voxels that contribute most strongly to a younger brain age prediction, orange ones the 5% contributing most strongly to an older prediction. 4b shows associations between voxel-wise brain-age scores and the whole-brain GM fraction for HPs and PwMS. In the upper row of 4b and superimposed on the axial and sagittal brain slices, we show significant voxel-wise t-statistics found for the association between voxel brain-age scores and GM fraction for HPs (orange palette) and for PwMS (blue palette) in independent analyses. The black arrowheads highlight the peak coordinate for which a significant negative overlapping association was found in independent analyses in both groups. The scatter graphs in the lower row of 4b illustrate the significant associations for this overlapping coordinate (i.e., at MNI: −47, −51, 44). 4c presents associations between voxel-wise brain-age scores and T2-weighted lesion load in patients, 4d illustrates group differences in these scores.

### Main Analysis 4: Functional connectivity and future brain-PAD in PwMS

This analysis showed that higher FC between right middle frontal cortex and cerebellar vermal lobule VI - VII during stress at the fMRI visit/baseline measurement in PwMS was associated with future brain-PAD increases or Δbrain-PAD respectively (t = 4.23, p_FWE_ = 0.0243; f^2^ = 2.98; Fig. 5).

**Figure 5.**
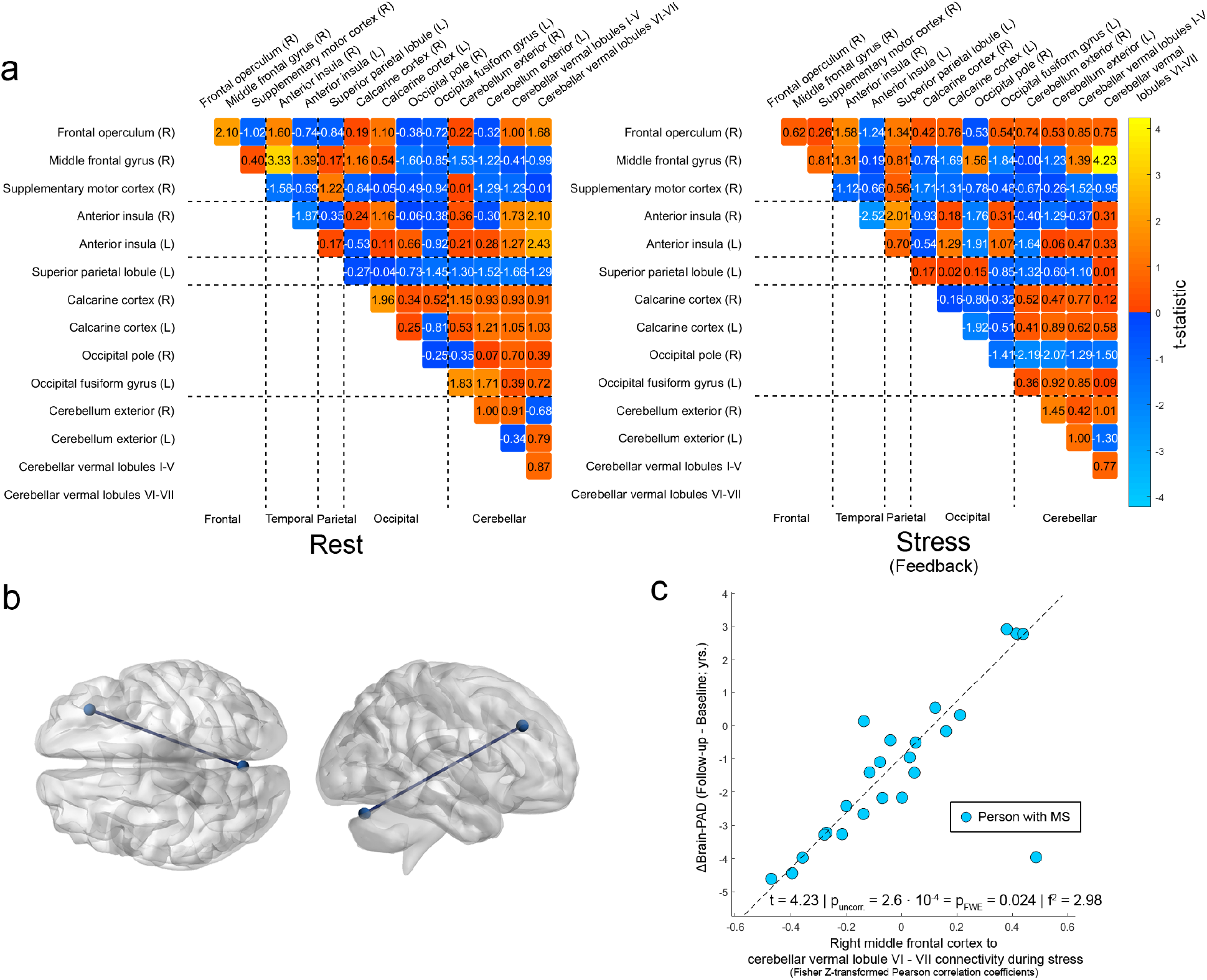
Functional connectivity and future brain-PAD in PwMS. The heatmaps in 5a depict the t-statistics for the association between FC at the baseline visit of pairs of stress-responsive regions and Δbrain-PAD on the left for rest and on the right for stress. 5b illustrates the connection between (the center coordinates) of right middle frontal cortex right and cerebellar vermal lobule VI – VII. 5c presents a scatter-diagram of this association.

## Discussion

Multiple findings suggest that psychological stress can impair brain health in health and disease alike. Consequently, we investigated whether comparable neurobiological stress brain-health pathways exist in health and MS with an fMRI stress task. We found that stress-stage FC between anterior insula and occipital regions was related to subclinical brain-PAD variations in HPs and to excessive brain-PAD in MS suggesting a generic neurobiological pathway linking stress and brain health.

Preparatory analyses showed that prerequisites for our main analyses were fulfilled. First, brain-age prediction accuracy (interpretable for this purpose in HPs only) was high: 81% of the variance in the chronological age of HPs was explained by predicted brain-age. Second, replicating (Cole et al., 2020) and (Kaufmann et al., 2019), patients’ brain-age was overestimated compared to HPs (6.1 years on average).

Main analysis one tested regional neural stress-responses within and between groups and showed that neural stress responses in both groups substantially overlapped with those observed in prior studies such as Dedovic et al. (2009) or Wang et al. (2005). Consistently, Supplementary analysis 1 showed that the task induced a stress-response in terms of heart rate and perceived stress in both groups. Crucially, however, none of the three measures differed between groups. Thus, despite findings of altered neuroendocrine stress processing in MS (Ysrraelitet al., 2008), the basic CNS, peripheral and psychological stress responsivity appears comparable in HPs and PwMS.

The second main analysis evaluated whether resting and stress-stage FC among stress-sensitive regions is related to brain-PAD independently for each group. As expected, only stress-stage FC was linked to brain-PAD. We found a positive association between FC of anterior insula and occipital areas with brain-PAD in both groups (HPs: right anterior insula - left occipital pole; PwMS: left anterior insula – left fusiform gyrus) which is consistent with the idea of generic neural pathway linking psychological stress to brain health. Noteworthy, at the same time, this idea is consistent with generic insular cortex functions. Anterior insula is a stress hormone sensitive (Serfling et al., 2019) hub region that measures stress-related peripheral inflammation (Slavich et al., 2010) and links these measurements to visual cognitive-affective processes (Sinha, 2014) such as threat cue monitoring performed in fusiform gyrus (Gomez et al., 2011). Importantly, Koren et al. (2021) could show that insula can not only measure an experimental peripheral inflammation but that reactivating the insular neurons active during inflammation at a later time can reactivate inflammation. Finally, cellular endpoints of the presumed generic stress – brain health pathway are suggested by work showing that immobilization stress reduces hippocampal neurogenesis (Chetty et al., 2014), induces irreversible dendrite loss (Radley et al., 2011) and increases Amyloid-β peptides (Ray et al., 2011) in healthy rodents.

Two supplementary analyses were conducted related to main analysis two that both aimed at evaluating the relative suitability of parameters selected for stress – brain health modeling in our study. Although again stress-stage anterior insula FC showed the strongest association to the brain health marker investigated in the first of the two (Supplementary analysis 2, GM fraction), the relations between FC and GM fraction were all weaker than those with brain-PAD and not significant on an FWE-corrected level. Supplementary analysis 3 tested associations between brain-PAD and markers of stress-response activity of regions in a neuroanatomical atlas but also failed to reveal significant associations. Together, these findings underscore the suitability of using stress-related FC assessed with ASL and brain-PAD for testing neurobiological stress – brain-health pathways.

Main analysis three evaluated within-group associations between regional brain aging and the whole-brain GM fraction to understand whether brain health as assessed by brain-age has a comparable neurobiological substrate in both groups. Compatible with Pagani et al. (2005) and Chen et al. (2004), parietal brain-age (i.e., in angular and supramarginal gyrus) showed a negative relation to patients’ GM fraction: The lower (higher) a patient’s GM fraction (atrophy), the higher these parietal coordinates’ brain-age. Importantly, a corresponding spatially overlapping association was found in HPs. These findings suggest that parietal over-aging is a generic marker of whole-brain GM loss and support the idea of a generic pathway connecting stress as starting and brain health as endpoint. Next, we related regional brain aging to the whole-brain volume of hyperintense lesions which revealed a strong positive association in periventricular WM – key areas of MS lesion occurrence (e.g., Polman et al., 2011). Moreover, we contrasted voxel-wise brain aging between groups. Consistent with thalamic MS atrophy (e.g., Azevedo et al., 2018), patients were characterized by elevated thalamic brain-age. Thus, the fact that the same single parameter – regional brain-age – is sensitive to indicators of the two major neuropathological MS phenomena – neuroinflammation and neurodegeneration – suggests that brain-age is a highly suited MRI marker of MS neuropathology. Moreover, even though several studies sought to separate MS neurodegeneration from aging-related neurodegeneration (e.g., Azevedo et al., 2019), this fact also suggests that MS neuropathology has (to a certain degree) an age-like profile.

Finally, in the fourth main analysis, we found that higher stress-stage FC between right middle frontal cortex and cerebellar vermal lobules is associated with higher future brain-PAD increases. This cinding shows that stress-related FC contains prognostic information for the future course of brain health in MS. More specifically, the fact that middle frontal gyrus is a key region for modulation of stress and emotions generated in other regions including the cerebellum (e.g., Ochsner et al., 2004) suggests that this prognostic link relates to prefrontal stress regulation.

One aspect of the study that could be discussed is that of using ASL for investigating FC. Although it is true that the majority of FC studies still uses BOLD MRI (Chen et al., 2015), ASL is a technique becoming increasingly popular for this purpose. Specifically, the review of Chen and colleagues from 2015 summarizes six ASL FC studies and especially in the clinical domain a variety of additional studies have been published since then. For example, Galazzo et al. (2019; on epilepsy), Boissoneault et al. (2016; Chronic Fatigue Syndrome), Liu et al., (2016; PTSD), and Fernandez-Seara et al. (2015; Parkinson’s Disease). Evaluating signal detection properties of ASL FC e.g., Vallée et al. (2020) found that by using standard ASL imaging protocols (i.e., 2D EPI pCASL sequence, the technique also used in our study), it was possible to detect established resting-state networks such as the default-mode and salience networks. Noteworthy, pCASL has excellent labeling efficiency and is thus the technique recommended for clinical applications by an ASL white paper (Alsop et al., 2014). Directly comparing ASL and BOLD-based FC, Jann et al. (2015) report that both techniques identified common resting-state functional networks which regionally overlapped on a moderate-to-high level. This study also found that reliability for networks identified with BOLD fMRI was higher than that for ASL-derived networks which still had a sufcicient reliability. This seeming disadvantage might, however, be more than compensated for by the higher robustness of ASL towards temporally slow signal artifacts (Wang et al., 2003; Aguirre et al., 2002). This is an especially important feature in a stress framework because these artifacts present in BOLD but not ASL can have a similar temporal profile as the slow (cf. Kirschbaum et al., 1993) stress response. Under these circumstances, artifact removal might be impossible unless one is not willing to remove the experimentally-induced signal (co-) variation simultaneously in BOLD fMRI. Consequently, given that our ASL acquisition parameters were well within the range of the abovementioned clinical ASL FC studies, given the exclusion of participants with strong head motion, the removal of residual motion artifacts and large-scale signal fluctuations during ASL preprocessing (Wang et al., 2012) described in the Supplement, using ASL for studying FC in a clinical and stress-framework is an appropriate choice.

Another point worth mentioning is that although the sample size was not predetermined by a sample size calculation, the study rests on a large to very large sample when compared to existing studies (i.e., larger than 88% of all MS fMRI studies employing cognitive tasks reported by Rocca et al. [2022] when only PwMS are considered, larger than 97% when also HP are considered).

A further important point is that of potential nuisance factors which pose a considerable challenge in any observational MS study due to its multifactorial pathophysiology and the large number of risk factors such as, but not limited to, Vitamin D level, dietary intake, or smoking (see e.g., Olsson et al., 2017 for an overview). We argue that potential nuisance effects were adequately accounted for here. For example, age, sex, severity of depressive symptoms, a measure of cognitive information processing, clinical disability, disease duration, presence of a progressive disease form, time-to-feedback, and study project (to control for the impact of slightly different anatomical MRI sequences employed in both projects) were modeled as nuisance factors with robust regression in main analysis 2. Importantly, the application of non-identical anatomical MRI sequences was not only accounted for by statistical modeling (a procedure also applied in multisite MS brain-age studies using different scanners such as the MAGNIMS study in which this factor accounted for 10.5% of the brain-PAD variation in patients; Cole et al., 2020) but also by the procedure employed by Bashyam et al. (2020) for training their brain-age classifier. In particular, their CNN was trained on a large heterogeneous collection of neuroimaging data comprising different data sets acquired with different scanners (14,468 individuals, 12 different datasets). Finally, however, the most important argument in favor of our assumption of a sufcicient nuisance factor consideration is the fact that stress-related anterior insular-occipital FC was linked to brain-PAD not only in PwMS but also in HPs in independent analyses. Due to this overlap, it is unlikely that disease-specific factors not considered were driving these associations in PwMS – because disease-related factors are not present in HPs.

A drawback of the study is that only patients of a single neurologic disorder were investigated, which limits the findings’ generalizability. Future studies should thus also include e.g., dementia patients despite their presumably more pronounced difficulties in participating in experimental settings given their comparably more severe average cognitive impairment (Hugo et al., 2014).

Taken together, we show that similar neural pathways link psychological stress and brain health in health and MS which points towards a generic stress – brain health pathway. The fact that brain-PAD is higher in MS despite these similarities suggests that the effect of this association is amplified by disease-specific vulnerability factors in MS. Finally, given findings that suggest that stress modification techniques can reduce brain-PAD in HPs (Luders et al., 2016) and lesions and neurodegeneration in MS (Burns et al., 2014; Mohr et al., 2012), our study could advocate the application of stress reduction techniques for promoting brain health independent of the presence of disease.

## Acknowledgements

We would like to thank all participants for taking part in this study and the Berlin Center for Advanced Neuroimaging for enabling the acquisition of MRI data. The work was supported by the German Research Foundation (WE 5967/2-1 and WE 5967/2-2 to MW, Exc 257 to FP, 389563835 and CRC 1404 – 414984028 to KR), the Brain & Behavior Research Foundation (NARSAD Young Investigator Grant to KR, USA) and the Deutsche Multiple Sklerose Gesellschaft (DMSG) eV (research award to KR). Our funding sources did not influence the study design, the collection, analysis and interpretation of data, the writing of the report or the decision to submit the article for publication.

