## Supplemental material for "Similar neural pathways link psychological stress and brain health in health and multiple sclerosis"

### SUPPLMENTARY MATERIAL

Short title: Neural stress processing and brain health

Key words: Multiple sclerosis, brain age, psychological stress, functional connectivity, machine learning, convolutional neural networks.

*These authors contributed equally

### Materials and Methods

#### Determination of heart rate

Pulse data during the rest and the stress fMRI measurement were measured with the standard pulse oxymeter of the Physiological Monitoring Unit provided with the MRI scanner. Because participating in the experimental paradigm included a manual operation of button boxes, the photoplethysmograph detector was attached to the participants' toes. The quality of heart rate signals measured in such a fashion has been proven to be comparable to that of signals assessed at participants' fingers (Hinkelbein et al. 2005).

For each participant and each of the two conditions, a single characteristic heart rate parameter was computed. The computation differed slightly across the two projects. In the first (see also Weygandt et al., 2016), we started by excluding heartbeats detected within the baseline pulse oxymeter raw signal (raw value signals below 2000) and those heartbeats that induced a heart rate acceleration to 133% or more or a deceleration to 75% or less. This was done because spurious pulse oxymeter signals may occasionally be detected as heartbeats which may falsely suggest a sudden increase of the heart rate, and because weak signals may lead to skipped heartbeats falsely suggesting an apparent decrease of the heart rate. The remaining signals during the full 8 minutes duration of the rest stage and the final 8 minutes of the stress stage were then used for heart rate averaging.

In the second (see also Brasanac et al., 2022), we started by computing a Fourier analysis based on the raw pulse signals of a given participant measured during the full eight minutes of the rest stage and one based on the raw signals measured during the final eight minutes (i.e., the feedback period) of the stress stage with the FFT algorithm implemented in Matlab. Frequencies below 30 Hz and above 150 Hz were discarded to constrain the fit to the physiologically meaningful frequency spectrum and amplitudes in two frequency windows (47.6 to 48.4 Hz, 95.6 to 96.4 Hz) were discarded due a technical artefact generated in these frequency ranges by the pulse oximeter. Finally, we fitted a unimodal gaussian with the Matlab FIT algorithm to the amplitude spectrum computed by FFT for both periods and took the frequency with the maximal fitted amplitude as single characteristic parameter for the given person and experimental stage

### MRI preprocessing

Preprocessing of anatomical brain scans and brain-age computation

Preprocessing of anatomical MRI images was performed using two similar pipelines, one for producing parameters needed in preprocessing of fMRI images ("Pipeline 1"), another for brain age computation ("Pipeline 2").

*Pipeline 1*

Basic preprocessing steps

A manual mapping of focal lesions in patients was performed by raters from the group of Prof. Paul first. This procedure was supervised by a neuroradiologist and was performed based on patients' T2 or FLAIR images. In a second step, probability tissue maps for GM, WM, and CSF were determined based on MP-RAGE images with the combined spatial normalization and segmentation algorithm implemented in SPM12. Voxel coordinates located in lesioned tissue as indicated by the lesion masks generated in step one (which were coregistered to the T1-weighted images beforehand) were omitted. This yielded voxel-wise tissue probability maps in the anatomical standard space defined by the MNI which were corrected for potential effects of local deformations applied during spatial normalization. Additionally, the lesion masks coregistered to the T1-weighted images were coregistered to the MNI space for computation of tissue-specific group mask (see below) using the spatial mapping parameters determined in the combined normalization and segmentation procedure.

Computation of GM fraction

The resulting voxel tissue probability maps were used to compute the GM fraction. Specifically, we first classified each voxel in MNI space as GM, WM of CSF by determining the maximal modulated tissue probability for a given voxel. If the coregistered lesion map contained a lesion at the voxel's coordinate, this voxel was always classified as lesion voxel. Afterwards, the GM tissue fraction for each participant was computed by dividing the number of GM voxels by the sum of all intracranial voxels.

Computation of group masks for GM, WM and CSF

Finally, group masks for GM, WM, and CSF were determined separately for PwMS in the cross-sectional study arm, for HPs in the cross-sectional study arm, and for PwMS in the longitudinal study arm. We computed the average tissue probability across all corresponding participants for each voxel and each of the three tissue types separately. Each voxel was then classified to the tissue for which the maximal average modulated probability was determined. Voxel coordinates located in lesioned tissue in at least one participant and the six voxels located in direct vicinity to such coordinates (i.e. with a Euclidean distance of no more than voxel) were removed from the group masks to account for partial voluming effects (e.g., Weygandt et al., 2011).

*Pipeline 2*

Basic preprocessing steps

In the second pipeline, we matched the less complex SPM12 normalization preprocessing applied by Bashyam et al. (2020), who provide a pre-trained convolutional neural network (CNN; https://github.com/vishnubashyam/DeepBrainNet) for brain-age computation. Specifically, we spatially normalized the T1-weighted anatomical scans and thus mapped them from participants' native to MNI-space with SPM12. Following Bashyam et al. (2020), however, this normalization exclusively used linear registration. Again, areas located in lesion tissue were omitted. Finally, we used a recently established method for skull-stripping of normalized T1-weighted scans which employs parallel two-dimensional U-Net-based CNNs and outperforms standard algorithms (CONSNet, Lucena et al., 2019). Linearly normalized and skull-stripped T1-weighted anatomical scans were then used for brain age computations.

<u>Preprocessing of functional brain scans</u>

ASL image preprocessing was performed with the ACID (Ruthotto et al., 2012) and the ASLtbx (Wang et al., 2008) toolboxes for SPM12. This comprised head motion correction, correction for distortions of the main magnetic field, and coregistration to the anatomical standard space defined by the MNI (voxel resolution $3 \cdot 3 \cdot 3$ mm$^3$) which utilized the spatial normalization parameters determined in preprocessing of T1-weighted images (see "Pipeline 1" above), and spatial smoothing. Subsequently, we extracted voxel-wise rCBF timeseries from the preprocessed ASL scans for both conditions (rest and stress) separately. All steps described here and below apply to the full eight minutes duration of the rest condition (stage 2) and the final eight minutes of the stress condition (stage 4). We focused on the final eight minutes of stage 4 as these were consistently included in the "Feedback" stress-substage in all participants.

**Framewise displacement-based fMRI motion assessment for participant exclusion**

A data quality assurance step was conducted based on the fMRI scans and using the framewise displacement (FWD) metric, an established fMRI quality assessment measure developed by Power et al. (2014) that evaluates the participants' fMRI head motion (determined during preprocessing of functional ASL scans) in a run-wise fashion. Within this run-wise approach, one average motion-based quality marker ("FWD-score") is determined for each condition (i.e., rest and stress) and participant.

After the FWD-score was determined for both conditions and all participants of each of the three datasets (i.e., for the 57 / 66 / 25 participants of Weygandt et al. [2016] / Brasanac et al. [2022] / Meyer-Arndt et al. [2020] who passed the clinicodemographic in- and exclusion criteria plus the visual quality assessment of anatomical images; see main text), we determined FWD outliers separately for each dataset and condition. Specifically, when an FWD-score in a condition of a dataset was more extreme than the first [third] quartile minus [plus] 1.5 times the Inter-Quartile Range of the FWD-scores computed for this condition and dataset, it was considered an outlier. Only those participants passed the procedure who had non-outlier FWD-scores in both conditions of a dataset. This was the case for 54 participants from Weygandt et al. (2016), 59 from Brasanac et al. (2022), and 24 from Meyer-Arndt et al. (2020).

Important to mention, the FWD-scores did not differ between PwMS and HPs. Specifically, we computed $t = 1.20$ ($p = 0.237$) in undirected two-sample t-tests for the FWD-scores of the rest measurement of the 57 participants in Weygandt, and $t = 0.71$ ($p = 0.478$) for the FWD-scores of the stress measurement. For the rest measurement of the 66 participants in Brasanc et al. (2022), we computed $t = -0.61$ ($p = 0.542$), and $t = -0.32$ ($p = 0.753$) for the stress measurement. Consequently, the abovementioned search for FWD outliers across data sets including PwMS as well as HPs appears legitimate.

**Within-participant regional stress responsivity**

To determine the regional neural stress responsivity, we first corrected the rCBF time series of each voxel, condition and participant that was (i) located in a region of the Neuromorphometrics brain atlas (http://Neuromorphometrics.com), (ii) covered by the GM mask of a given participant's group, and (iii) that contained non-zero rCBF for (i) the respective head motion parameters (Wang et al., 2012; Wang et al., 2008), and (ii) the global WM and CSF signals (Arnmann et al., 2015; Wang et al., 2012) of the given participant and condition using a linear model. Head motion parameters were determined by ASLtbx, the global WM and CSF signals were computed as the average signal across all voxels located in the group-specific WM and CSF masks.

After this correction, we computed an rCBF timeseries for each participant, condition, and region in the atlas that was averaged across timeseries of all voxels located in the region, covered by the group-specific GM mask, and that contained non-zero rCBF. If an atlas region did not include a single voxel fulfilling these criteria in one or more participants of a group, this region was excluded from the group's analysis. 121 of 122 atlas regions were included for HPs, 119 in PwMS. Left pallidum was not covered in HPs, left and right pallidum as well as cerebellar vermal lobules VIII-X were not covered in PwMS. This specific coverage pattern resulted from the relatively weak pallidal GM contrast in MP-RAGE scans (Droby et al., 2021), frequent MS lesions close to pallidum, and the proximity of cerebellar vermal lobules VIII-X to the inferior boundary of the field of view of the ASL sequence.

In the next step, we concatenated the average regional voxel time series of both conditions of a participant and computed region-wise stress responses with linear regression using the concatenated time series as dependent variable and a boxcar condition regressor coding zeros for the 60 rCBF scans included in the rest stage and ones for the 60 rCBF scans included in the final eight minutes of the stress stage as predictor (e.g., Wang et al., 2012; Wang et al., 2008). This was done for each participant and each region evaluated for the group of a given participant. The resulting region-wise regression coefficients were entered as regional stress-responsivity markers into main analysis 1.

**Statistical analyses**

Supplementary analysis 1: Psychological and physiological stress responses

Perceived stress and heart rate variations were evaluated based on the cross-sectional data with a factorial repeated measures analyses using linear mixed models (cf. Weygandt et al., 2019) testing main effects of condition, group membership, and their interaction on perceived stress and heart rate. For perceived stress (ratings during stages 3 & 5), data of 98 participants were available, for heart rate (average pulse across stage 2 & 4b) data of 84 participants. Each model included three fixed effects regressors (i.e., covariates of interest): One main effect regressor for experimental stage, one for group, and a regressor reflecting their interaction. Regressors for age, sex, project, cognitive task load, time to feedback, severity of depressive symptoms, plus constant modelled fixed effects of no interest. Clinical severity, disease duration, and progressive MS (y/n) were not modelled here as this might artificially reduce potential group differences as only patients have non-zero scores in these parameters. A constant modelling each participant's average perceived stress or pulse across experimental stages was included as random nuisance parameter. The type-I-error rate for the undirected tests performed were determined with permutation testing (10,000 within-subject permutations of each covariate of interest; cf. Winkler et al., 2014). An uncorrected significance threshold of $\alpha = 0.05$ was applied.

Supplementary analysis 2: Functional connectivity and grey matter fraction

This supplementary analysis was identical to the second main analysis except for the fact that we used participants' whole-brain GM fraction as dependent variable instead of brain-PAD.

Supplementary analysis 3: Regional neural stress response activity and brain-PAD

In this supplementary analysis, we used the regression coefficients computed for assessing regional neural stress responsivity in main analysis 1 to test these parameters' relation to brain-PAD. We applied an FWE-corrected significance threshold that corrected for the number of regions with significant stress responses (instead of the number of pairs of regions with significant stress responses) per group. Consequently, the uncorrected equivalent of an FWE-corrected threshold of $\alpha_{FWE} = 0.05$ corresponds to 0.05 / 17 = 0.003 in HPs and to 0.05 / 14 = 0.004 in PwMS. All other aspects were as described for the second main analysis.

### Results

#### Main Analysis 3: Group similarities and differences in regional brain aging

**Table S1** lists coordinates significantly related to overall grey matter volume and overall T2-weighted lesion load as well as coordinates separating significantly between groups. Abbreviations: CS, cluster size; Dist., distance.

| **Variable / Group / Region** | **Dist. (mm)** | **CS ($mm^3$)** | **x** | **y** | **z** | **t** | **$p_{FWE}$** |
|---|---|---|---|---|---|---|---|
| **Grey matter fraction** | | | | | | | |
| HP | | | | | | | |
| Supramarginal gyrus | 0 | 51 | -54 | -40 | 38 | -6.25 | 0.012 |
| Supramarginal gyrus | 2 | 20 | 56 | -22 | 32 | -6.25 | 0.013 |
| Supramarginal gyrus | 2.2 | 7 | -50 | -51 | 32 | -5.84 | 0.034 |
| Angular gyrus | 1 | 7 | -47 | -51 | 44 | -5.75 | 0.041 |
| Supramarginal gyrus | 0 | 7 | -39 | -48 | 44 | -5.73 | 0.043 |
| Supramarginal gyrus | 3.6 | 10 | -54 | -25 | 29 | -5.73 | 0.043 |
| Posterior cingulate gyrus | 0 | 3 | -9 | -46 | 2 | -5.67 | 0.048 |
| PwMS | | | | | | | |
| Angular gyrus | 0 | 2791 | -48 | -57 | 41 | -8.85 | $10^{-4}$ |
| Superior occipital gyrus | 0 | 128 | -26 | -90 | 19 | -6.33 | 0.003 |
| Middle temporal gyrus | 0 | 71 | -59 | -37 | -6 | -6.31 | 0.003 |
| Supramarginal gyrus | 0 | 128 | 54 | -34 | 55 | -6.24 | 0.004 |
| Supramarginal gyrus | 1.4 | 27 | -56 | -22 | 31 | -5.89 | 0.010 |
| Angular gyrus | 3 | 24 | -32 | -63 | 38 | -5.77 | 0.014 |
| Angular gyrus | 0 | 34 | 53 | -57 | 28 | -5.77 | 0.014 |
| Supramarginal gyrus | 2.2 | 17 | -42 | -49 | 41 | -5.68 | 0.018 |
| Supramarginal gyrus | 0 | 47 | -53 | -37 | 50 | -5.65 | 0.019 |
| Supramarginal_gyrus | 0 | 10 | 59 | -46 | 37 | -5.65 | 0.019 |
| Angular gyrus | 0 | 61 | 47 | -61 | 41 | -5.64 | 0.019 |

| | | | | | | | |
|---|---|---|---|---|---|---|---|
| Angular gyrus | 0 | 20 | -42 | -75 | 38 | -5.60 | 0.021 |
| Supramarginal gyrus | 1.4 | 7 | -45 | -31 | 35 | -5.36 | 0.042 |
| Angular gyrus | 0 | 3 | 44 | -70 | 37 | -5.34 | 0.043 |
| Middle frontal gyrus | 1.4 | 3 | -35 | 14 | 47 | -5.30 | 0.048 |
| Supramarginal gyrus | 0 | 3 | 63 | -36 | 25 | -5.28 | 0.049 |
| **T2-weighted lesion load** | | | | | | | |
| <u>PwMS</u> | | | | | | | |
| Cerebral white matter | 0 | 2028 | 33 | -49 | 10 | 8.45 | 0.0002 |
| Cerebral white matter | 0 | 135 | -23 | -25 | 26 | 7.00 | 0.0008 |
| Lateral ventricle | 0 | 361 | -26 | -51 | 8 | 6.64 | 0.0018 |
| Cerebral white matter | 0 | 41 | -27 | -51 | 22 | 6.60 | 0.0018 |
| Cerebral white matter | 0 | 159 | 21 | -22 | 28 | 6.51 | 0.0026 |
| Cerebral white matter | 0 | 138 | 20 | 35 | 8 | 6.19 | 0.0066 |
| Caudate | 0 | 78 | 17 | 23 | 2 | 6.18 | 0.0066 |
| Cerebral white matter | 0 | 54 | 14 | 29 | -7 | 6.15 | 0.0072 |
| Cerebral white matter | 0 | 44 | 42 | -54 | 11 | 6.14 | 0.0078 |
| Cerebellum exterior | 0 | 30 | -24 | -49 | -18 | 5.95 | 0.0146 |
| Cerebral white matter | 0 | 44 | -35 | -21 | 22 | 5.92 | 0.0160 |
| Cerebral white matter | 0 | 24 | -23 | -6 | 28 | 5.80 | 0.0228 |
| Cerebral white matter | 0 | 17 | 14 | 31 | 4 | 5.71 | 0.0294 |
| Cerebral white matter | 0 | 7 | -26 | 31 | 13 | 5.69 | 0.0324 |
| Cerebral white matter | 0 | 7 | -29 | -45 | 23 | 5.58 | 0.0406 |
| **Group** | | | | | | | |
| <u>HPs & PwMS</u> | | | | | | | |
| Thalamus proper | 0 | 213 | 18 | -27 | 11 | 5.51 | 0.0038 |

### Supplementary analysis 1: Psychological and physiological stress responses

Perceived stress and heart rate both increased significantly from rest to stress across members of both groups in the cross-sectional study arm (perceived stress: t = 12.31, p <$10^{-4}$, $f^2$ = 0.85; heart rate: t = 6.45, p <$10^{-4}$, $f^2$ = 0.46). Neither main effects of group, nor interaction effects were found. See Fig. S1 for details.

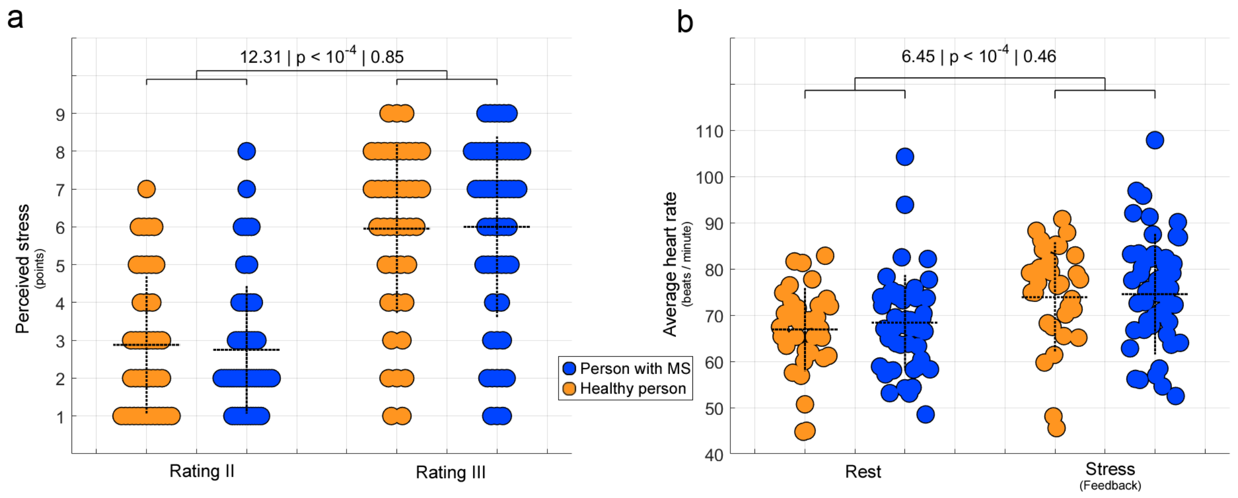

**Figure S1** illustrates the main effect of condition on perceived stress, 2e that on heart rate. The dotted horizontal lines depict the mean, the dotted vertical line the standard deviation for the respective parameter. The statistical parameters in 2d and 2e correspond to (from left to right) the t-statistic, p-value, and effect size $f^2$.

### Supplementary analysis 2: Functional connectivity and grey matter fraction

Testing associations between regional FC during rest and stress on one hand and GM fraction on the other separately for both groups did not reveal significant results on an FWE-corrected level. Worth mentioning, again the stress-related FC between right anterior insula and another region, i.e., inferior frontal gyrus, showed the strongest association between FC and GM fraction in healthy persons for all pairs of stress-responsive regions (t = -3.62, $p_{uncorr.}$ = 8.0 · $10^{-4}$, $p_{FWE}$ = 0.11, $f^2$ = 0.79). See Fig. S2 for details.

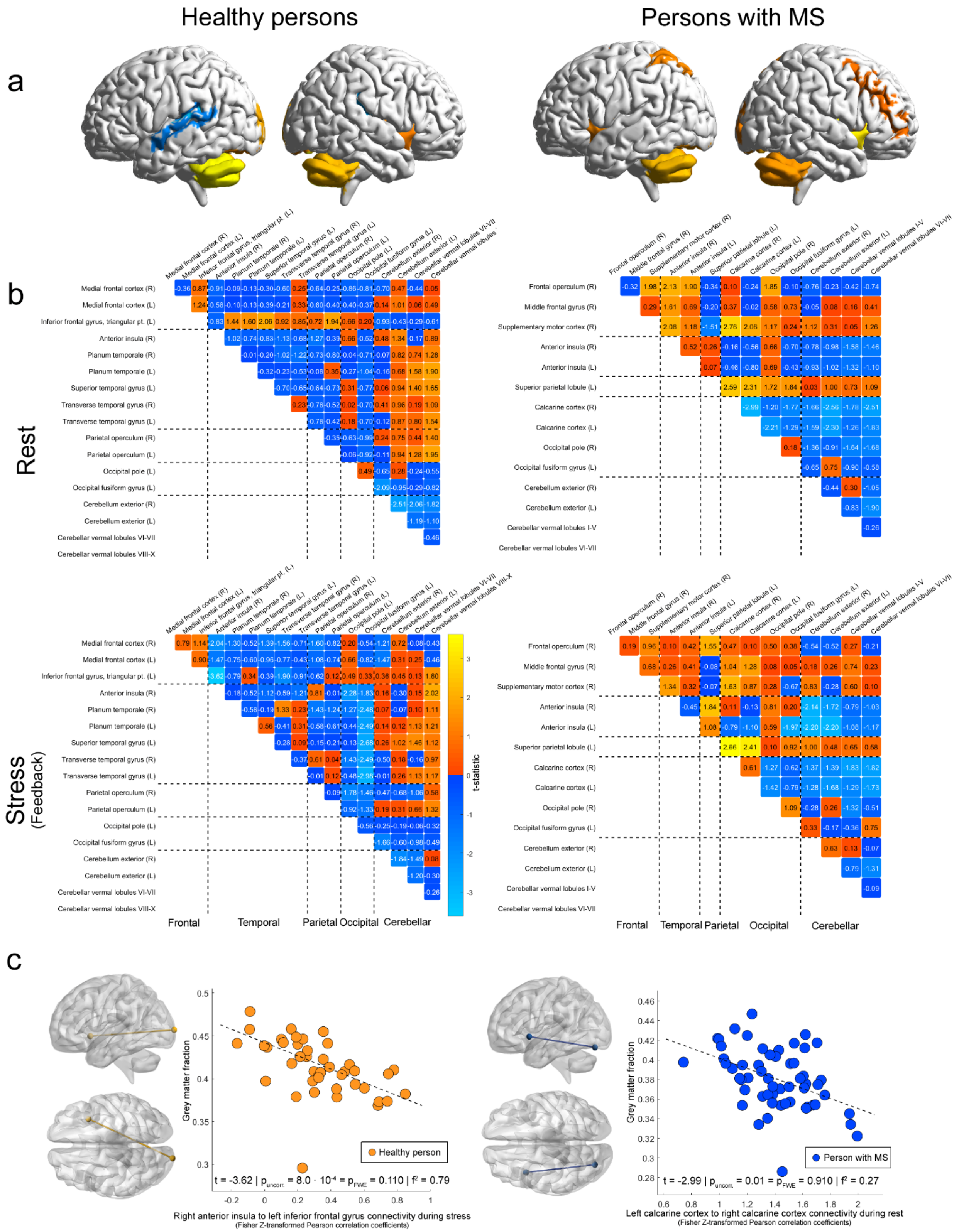

**Figure S2** illustrates associations between FC of stress-responsive brain regions and whole-brain GM fraction separated by group and condition.

**Supplementary analysis 3: Regional neural stress response activity and brain-PAD**

Regional neural stress responsivity was not related to brain-PAD – neither in PwMS nor in HPs. Fig. S3 below illustrates these findings.

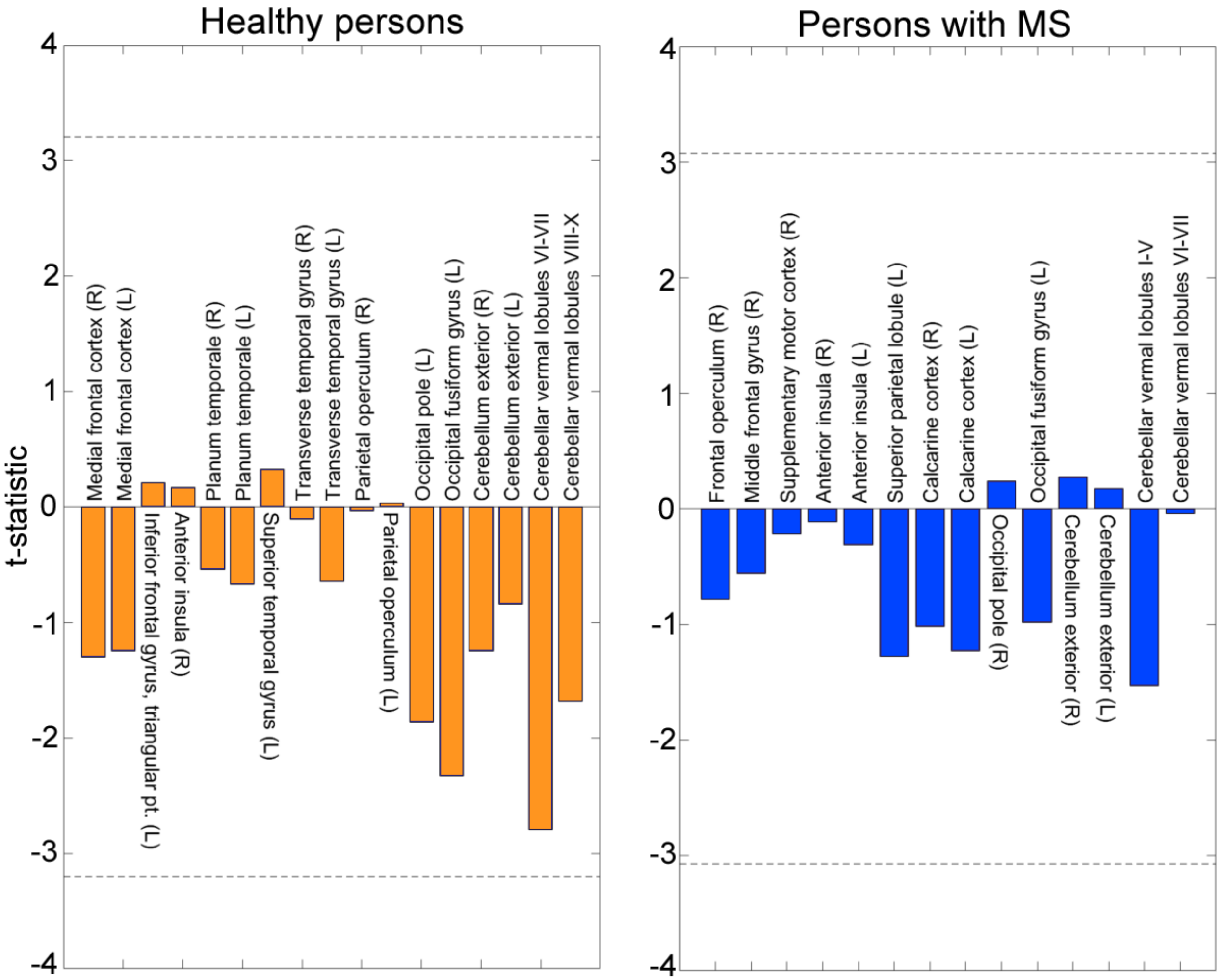

**Figure S3** illustrates associations between regional neural stress responsivity and brain-PAD separately for each group. The dashed line indicates the t-statistic necessary to achieve a significant undirected association on an FWE-corrected level.
